# Parallel genomic remodeling associated with independent terrestrialization events in arthropods

**DOI:** 10.64898/2025.12.29.696964

**Authors:** Lisandra Benítez-Álvarez, Vanina Tonzo, Leandro Aristide, Rosa Fernández

## Abstract

The repeated transition from aquatic to terrestrial environments (terrestrialization) has shaped the evolutionary trajectory of many animal lineages, yet the genomic basis of this ecological shift remains incompletely understood. Arthropods, with multiple independent terrestrialization events, provide a powerful system to investigate whether parallel genomic changes underlie adaptation to land. Here, we present a phylum-wide comparative phylogenomic analysis of 309 arthropod species representing aquatic and terrestrial lineages, using gene family evolutionary dynamics and directional selection analyses to uncover shared genomic strategies associated with life on land. We identified thousands of orthogroups showing parallel expansions or contractions across the three main lineages that colonized land (Arachnida, Myriapoda, and Hexapoda), of which a significant proportion also exhibited lineage-specific shifts in selective pressure. Functional enrichment of these orthogroups revealed functional convergence on processes such as oxidative stress response, transmembrane transport, energy metabolism, exoskeleton formation, and moulting regulation. Notably, parallel evolution in aquaporins, solute carriers, cytochrome P450s, superoxide dismutases, and heat shock proteins suggests that a conserved terrestrialization toolkit underlies independent colonization events. Additionally, parallel remodeling of key developmental and immune signaling pathways highlights the role of regulatory and structural innovations in adapting to terrestrial challenges. Our results provide the first large-scale genomic evidence of parallel molecular evolution driving arthropod terrestrialization and emphasize the power of comparative genomics to reveal shared solutions to ecological transitions across deep evolutionary timescales.

## Introduction

Animal terrestrialization, which involves the evolutionary transition from an aquatic to a terrestrial lifestyle, is undeniably one of the most extraordinary examples of life’s capacity to adapt and persist. This intricate process of evolution demands organisms to overcome important physiological and environmental obstacles, prompting the question of when, how, and how frequently species have successfully thrived in terrestrial environments (Dunlop et al., 2013). Adaptations of animals to terrestrial environments encompass a broad range of physiological processes and capabilities such as aerial respiration and gas exchange, water regulation and osmoregulation, thermoregulation, terrestrial locomotion, aerial sensory perception, reproduction outside of water including protection of eggs and embryos from water deprivation, utilization of novel food sources, and defense against environmental stresses like desiccation, rapid temperature changes, and heightened exposure to ultraviolet radiation (Frumkin & Chipman, 2023; Selden, 2005).

Multiple hypotheses exist regarding the environmental pathway that could have promoted the transition to terrestrial life. Among these, the main scenarios involve alternatively a marine–interstitial route (Little, 1990; Vermeij, 2020), a freshwater-to-terrestrial route (Glenner et al., 2006), and a subsurface route via caves (Frumkin & Chipman, 2023). These transitions were likely influenced by a combination of environmental pressures, such as changing water availability, oxygen levels, and competition for resources, which gradually drove adaptations that made survival on land possible.

By comparing the anatomical and physiological characteristics of aquatic and terrestrial animals, scientists have gained insights into the fundamental mechanisms that drive the evolutionary transition from sea to land. Prior research has emphasized the morphological and physiological changes that could aid in the transition to terrestrial life, including the development of tracheal systems (Hilken et al., 2021), as well as the presence of thickened cuticles, waxy coatings, spine-like outgrowths, sensory organs, and rigid exoskeletons (Asano et al., 2023). Despite these insights, many aspects of the transition to terrestrial life remain debated, including the relative influence of environmental pressures versus genetic predispositions in shaping these adaptations.

Metazoans have undergone multiple partial or full evolutionary transitions to terrestrial habitats in lineages such as in vertebrates (Ward et al., 2006); arthropods (Dunlop et al., 2013; Sharma, 2017); annelids (Struck et al., 2011); and mollusks (Aristide & Fernández, 2023; Kocot et al., 2011), among other phyla. The fossil record indicates that arthropods could be the earliest group of animals to inhabit land undergoing through at least three ancient (Hexapoda, Myriapoda, and Arachnida) and at least four more recent terrestrialization events (Isopoda, Amphipoda, Anomura, and Brachyura) (Rota-Stabelli et al., 2013). More specifically, evidence from fossils and trace fossils suggest that ancestral arthropods had the capability to venture into land as early as the Cambrian-Ordovician boundary, around 488 million years ago (Garwood, 2011), while myriapods and arachnids colonized terrestrial habitats during the Silurian period (ca. 416 to 443 million years ago; Ma). Shortly thereafter, during the Early Devonian (ca. 398-416 Ma), the fully terrestrial hexapods appeared. In addition, fossils from the Mesozoic period provide the earliest evidence of crustacean groups that lived on land. Molecular clock analyses, however, suggest that some of these groups may have emerged earlier than the fossil evidence indicates, with terrestrial myriapods, hexapods and arachnids purportedly appearing already during the Cambrian, the Ordovician, and the Ordovician-Silurian periods, respectively (Rehm et al., 2011; Wheat & Wahlberg, 2013). In particular, terrestrial arthropod species outnumber aquatic ones by a ratio of 17 to 1, indicating that the transition to land has been highly successful for this group, allowing them to diversify into an extraordinary array of ecological niches. This remarkable success can be attributed to key adaptations such as the development of waterproof exoskeletons, efficient tracheal respiratory systems, and specialized appendages for locomotion on land. Moreover, their ability to exploit novel terrestrial food sources, coupled with sophisticated reproductive strategies that minimize water dependency, has further facilitated their dominance in terrestrial ecosystems. As a result, arthropods have become the most speciose and widespread group of terrestrial animals, playing crucial roles in soil formation, nutrient cycling, pollination, and food webs across nearly all terrestrial habitats.

Several studies have been conducted to interrogate arthropod terrestrialization. These studies aimed to determine the timing and frequency of land colonization by different arthropod groups (eg, (Pisani et al., 2004; Rota-Stabelli et al., 2013; Sharma, 2017), as well as the factors contributing to the subsequent and significant diversification of terrestrial arthropods. Most studies on the subject primarily examine repeated phenotypic and physiological terrestrial adaptations within the main groups, possibly because of the importance of these traits to the success of life on land (Osorio & Bacon, 1994; Sharma, 2017). Nevertheless, the genomic basis for parallel adaptations among lineages on a large evolutionary scale, essential for comprehending both the origins of arthropods and their diversification on Earth, as well as the mechanisms that drive evolutionary adaptation, remains largely unknown.

This study seeks to bridge this knowledge gap by leveraging a phylogenomic approach to examine genome-wide gene repertoire evolution across the entire arthropod phylum. Specifically, we investigate how the interplay between gene family evolutionary dynamics and natural selection has contributed to the genomic reshaping of terrestrial arthropods. Our findings suggest that genome-wide parallel evolution, coupled with positive selection, has been a key driver in facilitating arthropod adaptation to terrestrial environments. These results provide new insights into the genetic mechanisms underpinning repeated evolutionary transitions to land and offer a broader perspective on how genomic innovations have shaped the remarkable success and diversification of terrestrial arthropods.

## Methods

### Taxon Sampling and Data Processing

A total of 656 arthropod species from all major lineages were chosen according to their availability in public databases to examine the genomic underpinnings of arthropod adaptation to terrestrial environments. In order to obtain the repertoire of coding proteins for each species, genomic data was filtered and processed as described in MATEdb (Fernández et al., 2022; Martínez-Redondo et al., 2024). In brief, fastp v.0.20.0 (S. Chen, 2023; S. Chen et al., 2018) was utilized to eliminate adapters and low-quality reads from the raw transcriptomic data. Trinity v.2.11 (Grabherr et al., 2011) was used with default settings to de-novo assemble data, and TransDecoder v5.5.0 (available at https://github.com/TransDecoder/TransDecoder) was used to predict candidate coding regions on the resulting assemblies. Then, contaminant sequences were filtered out using Blobtools 2 (Challis et al., 2020), and the longest isoforms of each transcript were retained for further analyses. When available, predicted proteomes were also directly downloaded from published arthropod genomes. The completeness of newly assembled transcriptomes and downloaded proteomes was assessed with BUSCO v.4.1.4 (Simão et al., 2015) in protein mode and the arthropoda_odb10 database. Only proteomes with BUSCO completeness scores greater than 85% (except for a few species with taxonomic importance with completeness values slightly lower than 85%) were included in subsequent analyses, resulting in 309 proteomes representatives of Arthropoda lineages, which were retained for further analysis. These covered arthropods of all lifestyles, to which we added two Onychophoran species as close outgroups and 10 species more as distant outgroups (see Supplementary Table S01). This broad species sampling is necessary to accurately identify changes in gene families that could be linked to habitat transitions. The main workflow including downstream analyses is described in Figure 1.

**FIGURE 1.**
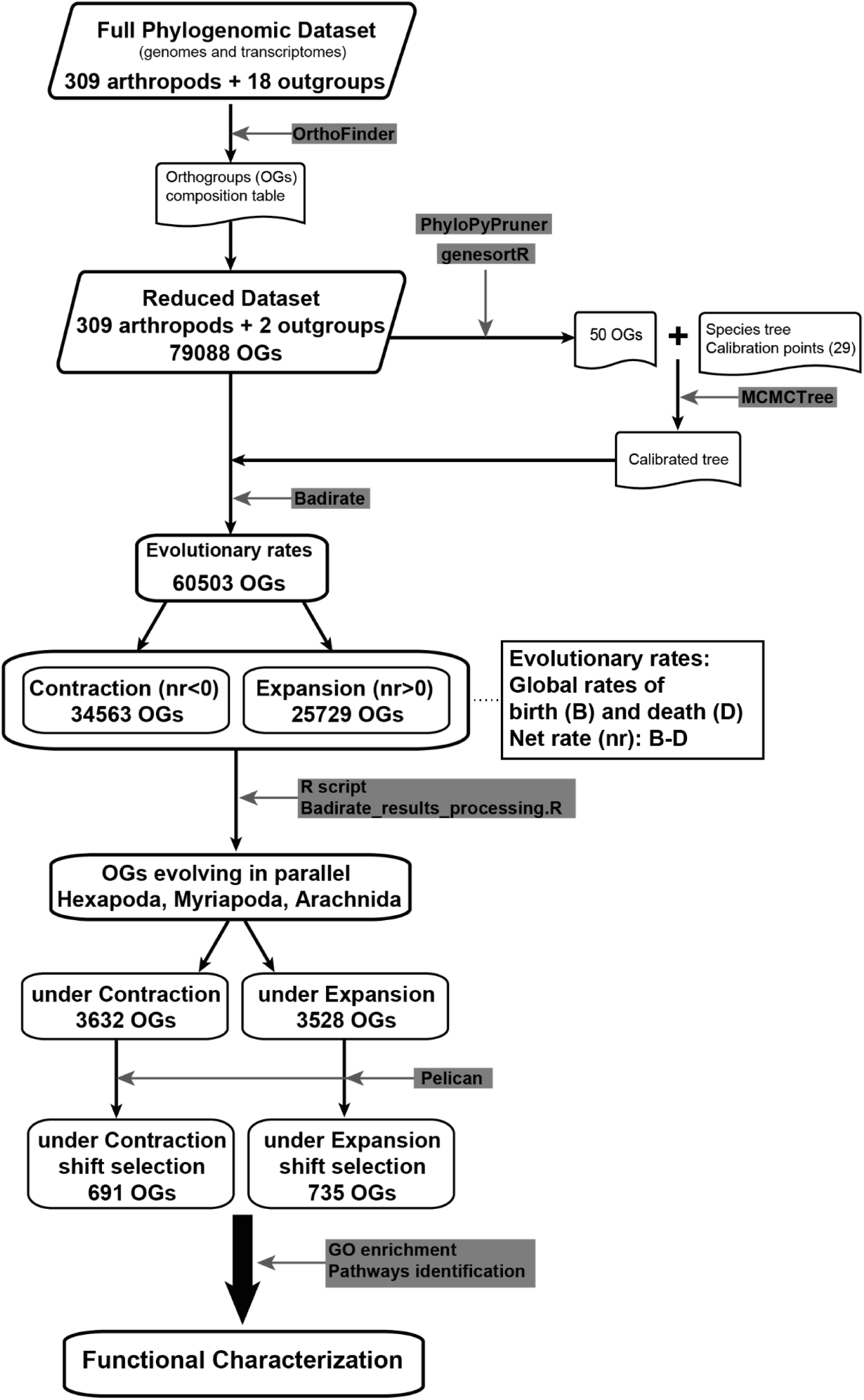
Workflow indicating the principal steps including in the methodology. Grey squares referred to the software used for each step.

### Orthology inference

We used OrthoFinder v.2.3.3 (Emms & Kelly, 2019) with default parameters to infer orthogroups in the final dataset composed of gene sets (protein mode) for 327 different species. Estimation of pairwise distances among proteome sequences was done with Diamond 2.0.8 (Buchfink et al., 2021) in sensitive mode. Orthology inference yielded 887,216 orthogroups (OGs hereafter), from which 79,088 had more than 4 sequences, which were retained for further analysis.

### Molecular dating

We manually obtained a tree topology for the sampled species by incorporating the most recent and accepted phylogenetic research on the phylum, plus the two onychophora species as closest outgroups (n=311 species; Supplementary Table S02, Supplementary Material 9). In particular, we adopted a topology consistent with the paraphyletic Arachnida hypothesis, which places Xiphosura nested within arachnids rather than as their sister group, following recent genome-scale phylogenomic studies employing stringent orthology filtering and site-heterogeneous models (e.g., Ballesteros & Sharma, 2019; Ontano et al., 2021; Ballesteros et al., 2022). Although alternative hypotheses exist, both alternative scenarios (a secondary aquatic return or multiple terrestrialization events within Arachnida) imply terrestrial ancestry for the group, and therefore are consistent with our interpretation of the genomic signatures linked to arthropod terrestrialization. This topology was leveraged to infer a time-calibrated phylogenetic tree using a subset of the orthologous genes inferred previously. To do so, we first masked and eliminated nonhomologous and low-quality sequence segments from each of the OGs with Prequal v1.02 (Whelan et al., 2018). Subsequently, masked OG sequences were aligned using MAFFT 7.4 (Katoh & Standley, 2013) with the automatic method selection. A phylogenetic tree was then generated for each of the aligned OGs, as follows. Trees for OGs with more than 500 sequences were inferred with FastTree 2.1.11 (Price et al., 2009) and the LG + CAT model as other methods were too computationally expensive for alignments of this magnitude. Trees for OGs with less than 500 sequences were inferred with IQ-TREE 2.1 (Minh et al., 2020) with the most suitable amino acid substitution model selected from around 95 different models, including mixture models (estimated in ModelFinder in the IQ-TREE 1.6.12 suite; (Kalyaanamoorthy et al., 2017). To minimize computation time, for the OGs with mixed models as the best-fitting, IQ-TREE was supplied with a guide tree inferred with FastTree. Ultrafast bootstrap support was calculated for all trees estimated by IQ-TREE using 1,000 iterations. Next, PhyloPyPruner (available at https://gitlab.com/fethalen/phylopypruner) was used to parse monophyletic groups of sequences within each OG (‘--prune LS’ option) option to extract the largest monophyletic subtree of orthologs per orthogroup, ensuring the retention of a single, well-supported orthologous sequence set per taxon while minimizing hidden paralogy. The following additional criteria were also enforced: (i) a minimum alignment length of 100 aminoacids (‘--min-len 100’), (ii) taxon occupancy of a minimum of 100 taxa (‘--min-taxa 100’), (iii) clades with a bootstrap value lower than 75 are discarded (‘--min-support 75’), (iv) for monophyletic groups of sequences belonging to the same species (i.e. in-paralogs), only the sequence with the shortest distance to the sister sequence of the group was retained (‘--mask pdist’). Although the minimum bootstrap threshold of 75 may appear stringent, it was deliberately chosen to prioritize alignment reliability and orthology confidence over taxon completeness, as low-support branches are more likely to include misassigned paralogs. After these steps, a total of 322 OGs that included a minimum of 80% of the species in the dataset were retained. The pruned OGs underwent additional analysis with geneSortR (available at https://github.com/mongiardino/genesortR) to rank them based on various statistics and features relevant for dating analyses, such as evolutionary rate, root-to-tip distance or saturation, among others. For molecular dating, we selected the 50 highest-ranking orthologs, which allowed us to minimise bias and reduce systematic error in subsequent analyses. Although this subset represents a small fraction of the available orthologs, molecular dating analyses with hundreds of taxa are computationally intensive, and increasing the number of loci dramatically inflates model complexity and runtime. Using the 50 top-ranked genes therefore represents a pragmatic compromise between computational feasibility and phylogenetic signal quality, ensuring stable and reliable divergence-time estimates while minimizing potential bias introduced by rate heterogeneity. This number strikes a good balance between information content and computational cost.

Finally, divergence time estimation analysis was performed using MCMCtree v4.9 (Rannala & Yang, 2007; Yang & Rannala, 2006) with an autocorrelated log-normal clock and using the approximate likelihood method (Reis & Yang, 2011), together with twenty-nine calibration points based on fossil data placed across the tree (Supplementary Table S03). Two parallel chains were executed for 600,000 generations, with a sampling frequency of 10,000. A burn-in period of 400,000 generations was applied, and the convergence of the chains was assessed by inspecting ESS values (>200) and posterior trace stability across independent runs.

### Reconstruction of the evolutionary dynamics of the gene repertoire of arthropods

We used the birth (B), death (D), and innovation (I) model implemented in BadiRate v1.35 (Librado et al., 2012; Librado & Rozas, 2022), to model and reconstruct the evolutionary dynamics of gene family size for each of the OGs across the arthropod tree. This maximum-likelihood-based approach estimates gene gains and losses for a set of OGs in a phylogenetic context. Specifically, we fitted a global rates (GR) model which assumes that all branches evolve under the same set of BDI rates. We used the OG counts per species table as inferred by OrthoFinder (Supplementary Material 1), excluding OGs with less than four sequences and the time-calibrated tree as input for the analysis. We ran BadiRate separately for each of the 79,088 OGs (i.e. we estimated OG-specific BDI rates) as very heterogeneous gene counts and the large dataset would cause BadiRate to crash when attempting to estimate only one set of BDI rates for the whole dataset. Even after this, for 21% of the OGs, the analysis failed to reach convergence during the likelihood calculation, and were discarded, finally obtaining results for a total of 60,503 OGs. The likely reason for this, after a careful exploration of the data, is the extremely skewed size distribution of these OGs (e.g. one or a few species with extremely large copy numbers) which may arise from idiosyncratic species-specific processes and factors that are most likely irrelevant for understanding large-scale, deep macroevolutionary patterns. Therefore, all conclusions taken from this study refer to ca. 80% of the gene repertoire (Supplementary Material 2). Following, based on the BDI model fit, we retrieved the minimum number of gene copy gains, losses, and the net size changes per branch for each OG (Supplementary Material 2). Against this background of OG expansions and contractions across the Arthropod tree we first aimed at identifying OG size changes phylogenetically associated with terrestrialization in the three oldest events: Arachnida, Myriapoda and Hexapoda, as these led to the most diverse groups and unique adaptations related to life on land. In this analysis, we classified all species using a binary terrestrial vs. aquatic scheme, consistent with previous large-scale comparative frameworks for studying habitat transitions across metazoans (Rota-Stabelli et al., 2013; Aristide & Fernández, 2023; Martínez-Redondo et al., 2023). This coarse-grained categorization reflects the deep evolutionary timescales of the terrestrialization events considered here, where finer ecological distinctions such as semi-terrestrial, amphibious, or microhabitat-level differences cannot be reconstructed with confidence. Furthermore, the focal terrestrialization events leading to Arachnida, Myriapoda, and Hexapoda represent ancient, well-established ecological transitions that have remained stable within these lineages (Dunlop et al., 2013; Sharma, 2017). Therefore, this binary classification accurately captures the main ecological contrast relevant to our evolutionary framework. Then, we looked for parallel evolutionary dynamics in these OGs across the three lineages, as parallelism could be a strong signal of the process of adaptation to the terrestrial environment. We set up two criteria to be fulfilled for establishing parallelism: (i) that the direction of size change (contraction or expansion) of a given OG in the considered terrestrial lineages be the same (e.g. if an OG had net gains in each of the three terrestrial lineages, it was considered an OG under parallel expansion); (ii) that these parallelly expanded/contracted OGs exhibited shifts in selective pressure in the terrestrial branches compared to non-terrestrial ones, linking the adaptive terrestrialization processes to the previously identified parallel evolution of the OGs (see next section, Supplementary Material 3). To clarify, our criterion for identifying parallel changes focused on the directionality of gene family evolution rather than rate acceleration: we required the same net change (expansion or contraction) to occur independently in all focal terrestrial lineages and in no other branches of the tree. This approach aims to capture consistent genomic responses to terrestrial adaptation, which may be biologically meaningful even when evolutionary rates are not exceptional, while remaining stringent enough to minimize false positives.

### Inference of selective shifts in OGs evolving in parallel

To determine if the OGs that evolved in parallel in terrestrial lineages underwent shifts in selection regime, which could indicate adaptive changes associated with the transition to land, we used Pelican v.1.0.8. (Duchemin et al., 2023). This program utilizes amino acid profiles to identify shifts in selection direction (Parto & Lartillot, 2018) and pinpoint sites that are associated with a trait change throughout the phylogeny. Pelican examines the hypothesis that the substitution process at each site in a protein sequence is contingent upon a given trait or environmental condition. To accomplish this, the likelihood of an homogeneous model where the substitution process is identical across the whole tree is compared, using a likelihood ratio test (LTR), to a condition-dependent model where terrestrial and non-terrestrial branches of the tree evolve under different substitution processes. Treerecs v.1.2 (Comte et al., 2020) was used to reconcile the gene trees corresponding to the OGs included with the species tree that had been previously employed by MCMCtree for time tree inference. We coded non-terrestrial as background and terrestrial as foreground characters, respectively. Finally, we ran Pelican on the 6100 OGs previously identified as having parallel size changes using the reconstructed gene tree for each of these OGs, annotating the branches as background (non-terrestrial) or foreground (terrestrial), as required by Pelican, following ancestral state reconstructions under the maximum parsimony algorithm using a custom R script (Supplementary Material 4).

Finally, to identify the OGs that exhibit a significant shift in the selection regime associated with the terrestrial condition, we aggregated site-level p-values for each OG to obtain an OG-level p-value using the Gene-wise Truncated Fisher’s (GTF) approach (Douchemin pers. comm.; see Supplementary Material 5).

### Functional annotation, gene ontology enrichment, and functional similarity analysis

For all the species included in the analysis, we functionally annotated the amino acid sequences of each proteome using eggNOG-mapper v.2.1.6 (Cantalapiedra et al., 2021). We used the DIAMOND algorithm for the search step using the Arthropoda database, and produced OG-level annotations by extracting gene ontology (GO) terms from each sequence within each OG and filtering unique GO terms. To further investigate the putative functions of the parallelly-evolving OGs with shifts in directional selection (both expanded and contracted), we performed overrepresentation tests using the R package clusterProfiler v.3.10.1 (Yu et al., 2012) (*P* < 0.05), using as background set the combination of all unique GOs for all OGs (Supplementary Material 7 and 8 for expanded and contracted OGs, respectively). The results were visualized with the rrvgo v.1.14.0. R package (Sayols, 2023), using the *Drosophila melanogaster* database and the Wang method to estimate the semantic similarity among GO terms (see GOoverrepresentation_visualization.R in Supplementary Material 6).

To further investigate the pathways in which the parallel evolving OGs under shifts in selection were involved, we performed pathway enrichment with PANGEA (Hu et al., 2023) (available at https://www.flyrnai.org/tools/pangea/web/home/7227), using the FlyBase signaling pathways as the reference, which is based on experimental evidence.

## Results

### Gene repertoire evolutionary dynamics across Arthropoda

Gene repertoire dynamics was inferred implementing a phylogenetic model that incorporates a stochastic process of gene family size evolution to identify changes in the gene repertoire that could potentially be associated with the process of transition to life on land (Table S04, Supplementary Material 2). We identified 25,729 OGs exhibiting a consistent trend of expansion along the phylogeny, as indicated by a positive difference between birth and death rates, while 34,563 OGs experienced contraction, indicated by a negative difference. In this context, gene family expansions or contractions refer to changes occurring within individual lineages across the tree, rather than to patterns observed globally across all taxa. The reconstruction of the global gene repertoire dynamics (i.e. number of gene copy gains and losses in each branch; Figure 2) indicated that deeper nodes have a high rate of gene loss. On the other hand, gene family expansions (OGs with high copy number) appear to be predominant in shallower branches of the tree, revealing a highly dynamic repertoire evolution across arthropods in which habitat transitions do not seem at first glance to be the main driver of repertoire changes (Fig. 2).

**FIGURE 2.**
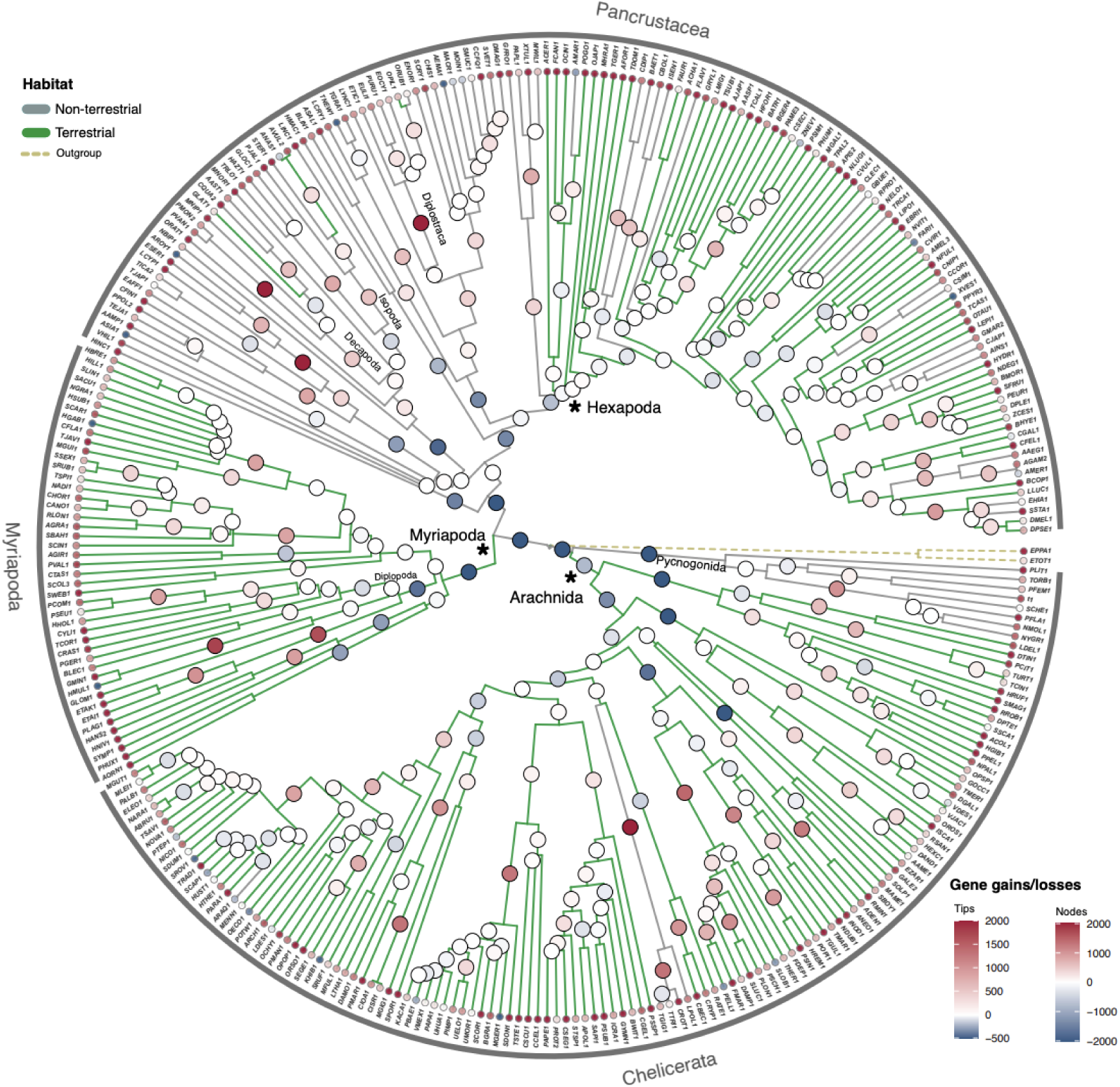
Arthropoda time tree with mapped net gains and losses obtained from Badirate using a Birth-Death-Innovation (BDI) model with all branches evolving under the same set of BDI rates. The circles represent the gains and losses on a different scale for nodes and tips, as shown in the legend. The gradient of colour represents the amount of gains (red) and losses (blue). The habitat of the branches is indicated in gray (non-terrestrial) and green (terrestrial).

### Parallel evolving gene families under directional selection in terrestrial lineages

After calculating the net gains and losses per branch for each OG from the Badirate analyses, we observed that Myriapoda and Arachnida (the first ones transitioning to land) have a larger number of OGs with expansions (17,688 and 25,036 OGs, respectively) and contractions (11,938 and 20,990 OGs), compared to Hexapoda (15,462 and 8,188 OGs under expansion and contraction, respectively) (Fig. 3). Overall, there is a trend for expansion to be the main characteristic of the evolution of the OGs analyzed in the three lineages (Figure 3). We next tested whether there was parallel expansion and contraction among these three main terrestrialized lineages. We recovered a total of 3528 and 3622 OGs under parallel expansion and contractions, respectively (Figure 3).

**FIGURE 3.**
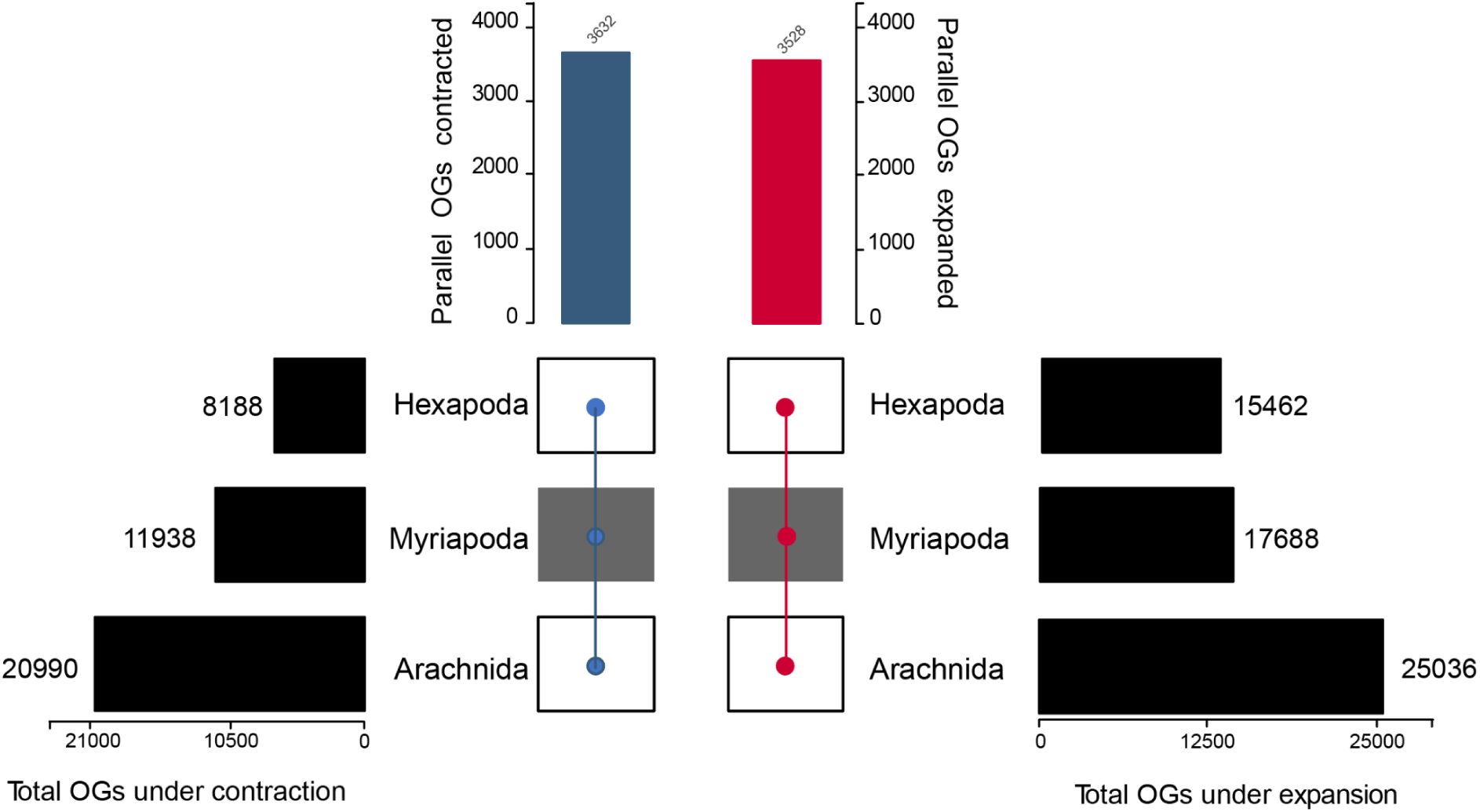
Parallel expansions and contractions across terrestrial lineages. The number of orthogroups (OGs) under contraction (left) and expansion (right) are shown for each one of the three terrestrial lineages (Arachnida, Myriapoda, and Hexapoda). The total number of parallel evolving OGs for all these lineages is indicated in a bar chart on the top.

To understand the adaptive potential of these parallel evolving OG expansions and contractions related to terrestrial habitats, we examined if these were further subjected to shifts in directional selection associated with the habitat change in the focal lineages. Out of the 3,528 OGs that showed parallel expansion, a total of 2,330 OGs could be analyzed with Pelican (the remaining failing potentially due to insufficient sequence conservation, low alignment quality, or missing data in key taxa). From these, 735 OGs showed a significant shift in directional selection regime (∼32% of parallelly expanded OGs; Supplementary Table S05). Similarly, out of the 3,632 OGs that experienced parallel contractions, 2,366 OGs could be further analyzed with Pelican, out of which 691 OGs showed evidence of selective pressure changes (∼29% of parallelly contracted OGs; Supplementary Table S06). These results unveil a high number of OGs evolving in parallel with convergent shifts in directional selection.

### Functional characterization of parallel evolving gene families across terrestrial arthropods under shifts in directional selection

In order to identify genes potentially involved in adaptation to life on land across the three arthropod lineages, we next performed a functional characterization and enrichment analysis of GO terms annotated to the OGs under parallel size changes and shifts in directional selection. While GO-term enrichment provides a useful overview of functional trends, we recognize that individual GO annotations, particularly in non-model taxa, can be overly specific or biologically misleading due to the propagation of functional labels from distantly related model organisms. Therefore, the patterns described here should be interpreted as high-level functional signatures rather than literal mechanistic inferences. Our analysis aims to highlight broad, recurrent categories (e.g., metabolism, regulation, stress response) that emerge consistently across independent terrestrial lineages, serving as a starting point for more targeted, gene-level functional studies as genomic annotations improve.

Enrichment analysis of OG parallel expansions (Figure 4) revealed key functional categories associated with metabolic processes, cellular regulation, and organismal adaptation. Notably, we observed significant overrepresentation of energy metabolism pathways, including ATP metabolic processes, mitochondrial respiratory chain complex I assembly, and cellular respiration, which may reflect adaptations to terrestrial environments. Carbohydrate and amino acid metabolism, alongside protein biosynthetic processes, were also enriched, indicating potential shifts in nutrient processing. Several categories linked to cellular regulation, such as transcriptional control, nucleotide biosynthesis, and gene silencing by RNA, highlight mechanisms of regulatory adaptation. Additionally, processes related to cellular stress responses, detoxification, and metal ion sequestration suggest the importance of homeostatic regulation. Notably, the expansion of oxidative stress response genes and heat shock proteins (HSPs) could enlarge the cellular protection mechanisms. Interestingly, enriched terms related to inter-male aggression, muscle differentiation, and cloaca development may indicate lineage-specific adaptations in behavior and reproductive biology.

**FIGURE 4.**
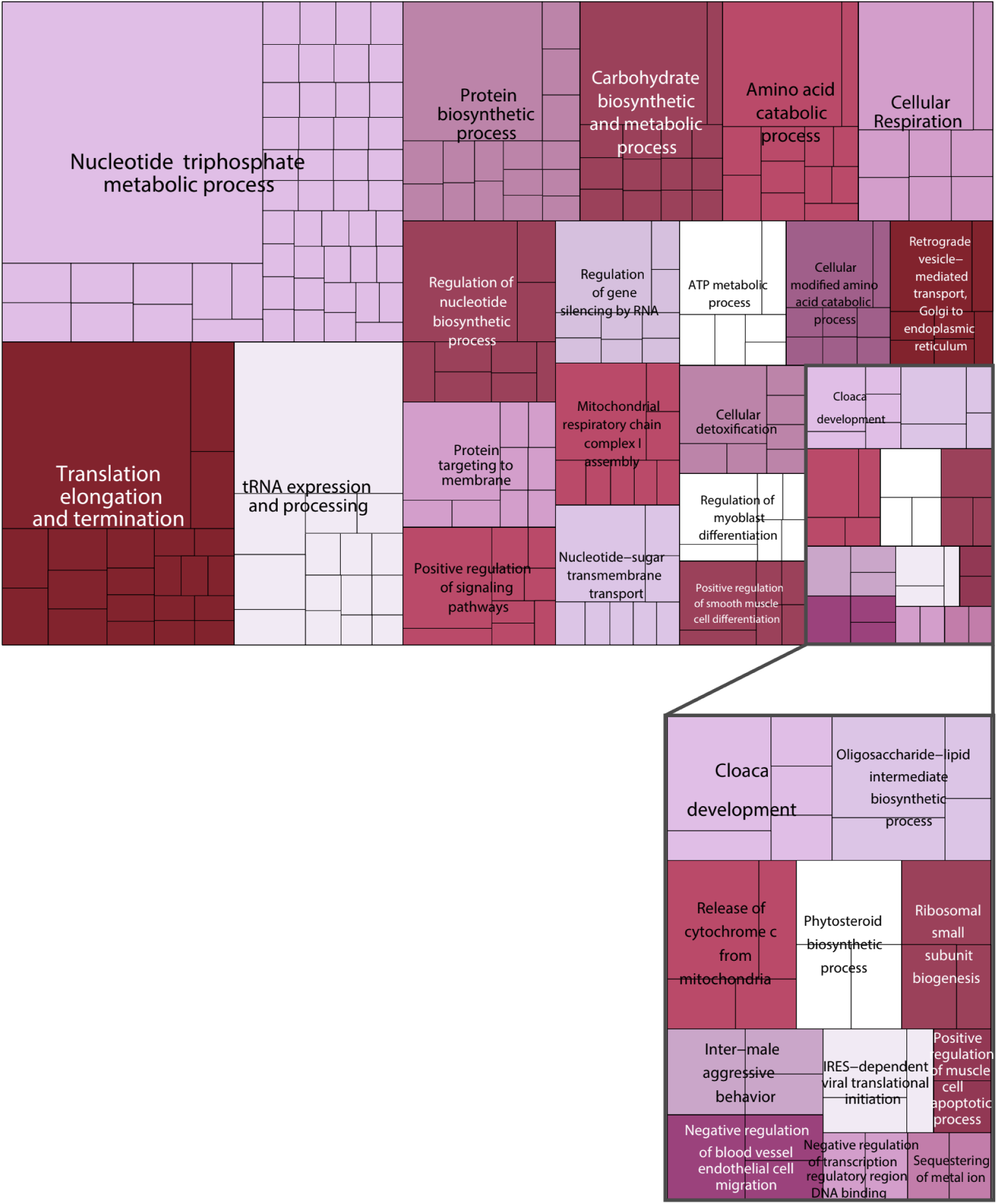
Treemap showing overrepresented GO terms (Biological Process) of parallel evolving OGs under expansion and with significant directional selection shifts Myriapoda, Hexapoda and Arachnida.

Enriched GO terms in OGs under parallel contraction and a shift in directional selection in the three terrestrial lineages (Figure 5) revealed a loss or reduction of functions related to protein homeostasis, metabolism, and cellular regulation. Notably, several categories involved in mitochondrial function, oxidative phosphorylation, and ATP metabolism were enriched, suggesting potential shifts in energy production strategies. Processes related to RNA metabolism, translation, and ribosomal biogenesis were also contracted, indicating possible streamlining of gene expression regulation. Additionally, we observed significant reductions in pathways associated with lipid metabolism as well as protein folding, and endoplasmic reticulum stress responses, hinting at metabolic stress response. Several immune-related pathways, including negative regulation of viral-induced signaling and cytoplasmic pattern recognition receptor activity, suggest shifts in host-pathogen interactions. The enrichment of terms related to apoptosis, neurodevelopment, and muscle cell differentiation could indicate trade-offs between cellular maintenance and developmental plasticity. Interestingly, reductions in epithelial cilium movement, epithelial polarity maintenance, and tracheal system branching suggest potential morphological adaptations. Collectively, these contractions highlight functional shifts that may reflect both energetic trade-offs and adaptive modifications in arthropod terrestrialization.

**FIGURE 5.**
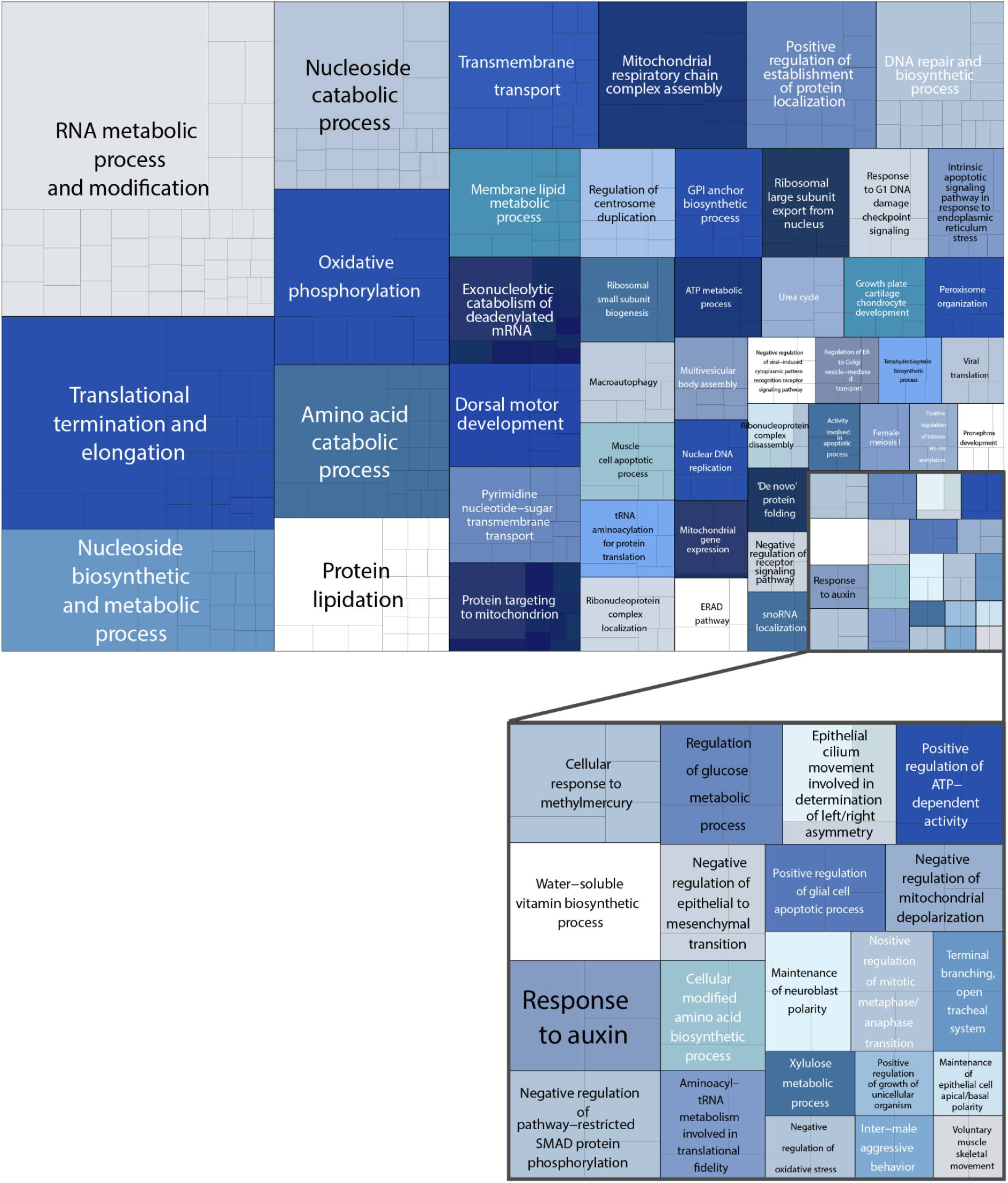
Treemap showing overrepresented GO terms (Biological Process) of parallel evolving OGs under contraction and significant directional selection shift in Myriapoda, Hexapoda and Arachnida.

The enrichment patterns for parallel expansions and contractions show overlap in several functional categories, particularly in metabolism, gene regulation, and development. Both expansions and contractions include terms related to ATP metabolism, mitochondrial function, and oxidative processes, as well as gene expression, RNA metabolism, and protein biosynthesis. Developmental pathways appear in both datasets, with terms linked to muscle differentiation, apoptosis, and cellular organization. Additionally, categories related to transcriptional regulation, nucleotide biosynthesis, and metabolic processes are present in both sets of enrichments. These shared functional themes suggest that similar biological processes were affected by both expansions and contractions, potentially reflecting a general reshaping of the genetic machinery underlying the biological systems involved in these processes, driven by common adaptation to the new terrestrial environments.

To find the pathways that the parallel evolving OGs are involved in (regardless of whether they are contracting or expanding), PANGEA used the FlyBase gene set to find pathways that have been curated by experimental evidence in D. melanogaster (Supplementary Table S7). These pathways can be grouped into 5 processes: metabolism and reproduction, immunity, embryogenesis, development, and cell differentiation and morphogenesis (Table 1, Supplementary Table S7, S8).

**TABLE 1.**
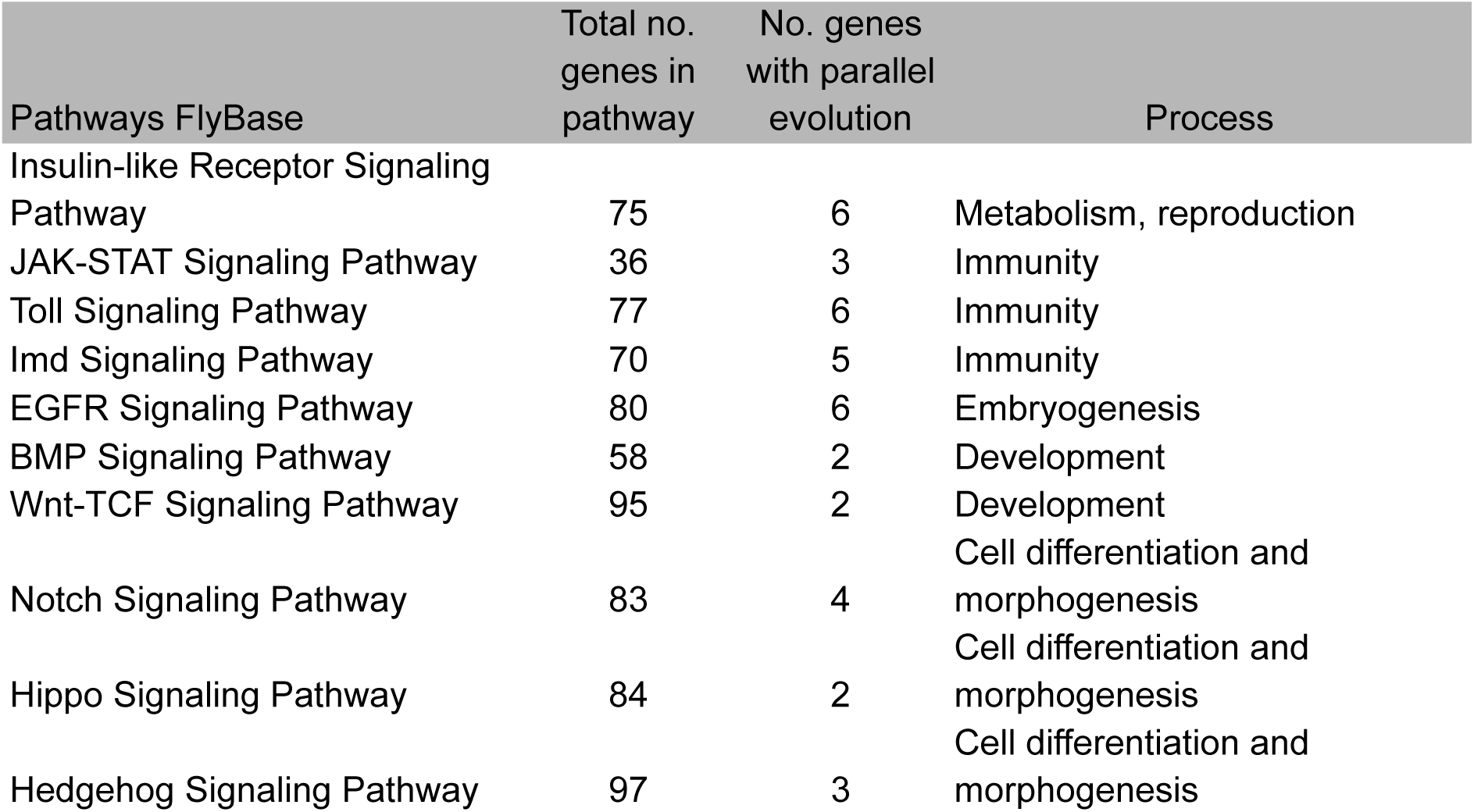
Pathways in which the parallelly-evolving OGs under selection are involved. Total number of genes included in each pathway as well as the number of genes with parallel evolution identified here are indicated.

The identified pathways involved in metabolism, immunity, development, and cellular regulation, with several key signaling pathways in FlyBase showing potential links to these changes. The Insulin-like receptor signaling pathway, which regulates metabolism and energy balance, aligns with the expansion of ATP metabolism and mitochondrial processes, suggesting metabolic adaptations to terrestrial environments. Immune-related pathways, including JAK-STAT, Toll, and Imd signaling, correspond to contractions in viral recognition and immune regulation, indicating potential shifts in pathogen defense strategies. Developmental pathways such as EGFR, BMP, and Wnt-TCF signaling, which control cell differentiation and tissue growth, relate to expansions in muscle differentiation and apoptosis regulation, as well as contractions in epithelial organization and cilium movement, suggesting morphological changes. The Notch and Hippo pathways, involved in cell fate determination and growth regulation, connect to expansions in differentiation control and contractions in polarity maintenance, pointing to shifts in developmental regulation. Finally, the Hedgehog pathway, which influences pattern formation, may be linked to observed changes in organogenesis and tissue organization. These results suggest that these key signaling networks played a role in both adaptive expansions and selective contractions during arthropod terrestrialization. While these pathway-level enrichments provide an informative overview of the main biological systems affected, we note that some broad categories—such as immunity or development—encompass highly diverse gene families whose functions may vary across arthropod lineages. Therefore, these results should be interpreted as reflecting general functional trends rather than precise mechanistic pathways, as part of the observed signal may also derive from lineage-specific morphological or ecological diversification unrelated to terrestrialization.

## Discussion

### Parallel genomic evolution driving arthropod terrestrialisation

Gene family expansion and contraction has been associated with ecological habitat shifts and adaptive success in different arthropod lineages (Balart-García et al., 2023; Chattopadhyay et al., 2020; Vertacnik et al., 2021). Our results emphasize the critical role that gene family expansions and contractions play in the recurring shift to terrestrial life across the three main terrestrial arthropod lineages. The detection of OGs evolving in parallel with convergent shifts in directional selection suggests that arthropods have followed common genetic and evolutionary paths towards their adaptation to land. In this sense, our results support the idea that important innovations arose via parallel evolution, helping these lineages adapt to life on land. Among the most enriched GO categories during parallel expansion followed by shifts in directional selection, we identified stress-response systems, cellular control, and metabolic pathways. Expansions in ATP metabolism, mitochondrial respiratory chain complex I assembly, glucose and amino acid metabolism suggest a necessary crucial improvement in metabolic efficiency and energy output needed for terrestrial adaptation. This is consistent with previous studies showing increased metabolic rates associated with the success of terrestrial arthropods (Harrison et al., 2010; Misof et al., 2014). Moreover, the expansion of oxidative stress response genes and heat shock proteins points to enhanced cellular protection mechanisms potentially helping arthropods resist some of the challenges associated with life on land, such as desiccation, temperature fluctuations, and UV exposure (Benoit et al., 2023; Klok et al., 2004; Lucu & Turner, 2024). It is unlikely that these recurrent expansions arose by chance, as our inference of gene family dynamics relied on explicit birth–death modeling across a fixed, time-calibrated phylogeny, with duplication and loss rates statistically estimated for each branch. Moreover, only gene families showing the same direction of change across all three terrestrialization branches, and in none of the others, were retained, a conservative criterion that strongly reduces the probability of stochastic coincidences. Importantly, independent studies based on different methodological frameworks have reported comparable patterns; for example, Martínez-Redondo et al. (2023) detected parallel expansions of aquaporin gene families in terrestrial arthropods using detailed gene-tree reconciliation and molecular evolution analyses, providing further support that these signatures represent genuine adaptive responses rather than random effects.

In contrast, parallel contractions in terrestrial arthropod lineages were mostly related to energy control, immune system, and developmental pathways. The loss of gene copies linked to oxidative phosphorylation, ribosome biogenesis, and mitochondrial function indicates that whereas certain metabolic processes expanded, others were streamlined, potentially reflecting energetic trade-offs required for life on land. In particular, the contraction of immune-related pathways, especially those linked to viral recognition and pattern recognition receptor signalling, may point to changes in host-pathogen interactions, potentially resulting from a reduced pathogen diversity in terrestrial habitats (St Leger, 2021). These results underscore the complex balance between gene expansion and contraction in shaping evolutionary responses to new ecological pressures.

### Towards the identification of an arthropod terrestrialization genomic toolkit

Our results suggest that a core genomic toolkit—shared across major terrestrial arthropod lineages—has evolved under positive selection during terrestrialization. This toolkit includes, among others, gene families involved in transmembrane transport. Several OGs annotated as aquaporins (e.g., AQP3, AQP9) and ATP-dependent ion transporters were found under parallel expansion and selection. Aquaporins facilitate water and small solute movement and are central to osmoregulation, a crucial function in terrestrial settings (Aristide & Fernández, 2023; W. Wang et al., 2023). Aquaporin evolution is also tied to terrestrialization in mollusks (Martínez-Redondo et al., 2023) and amphibious fish (Lorente-Martínez et al., 2023).

Gene families related to exoskeleton formation and moulting also showed signs of selection and expansion, including chitin synthase (CHS), Notch, PTTH, and various cytochrome P450s (e.g., cyp307a1, cyp314a1). The exoskeleton provides desiccation resistance, structural support, and enables morphogenetic flexibility, and its evolution is key to terrestrial success in arthropod, including complex behaviours related to matting courtship or territory defending (Asano et al., 2023; Belles, 2019; Ito & Uchiumi, 2024; Lehmann et al., 2024).

The HSP70, DNAJC9, and STI1 gene families, crucial in protein folding and repair, show expansions with selection shifts, underlining their role in thermal and oxidative stress responses. HSPs are central to environmental resilience, with inducible and constitutive forms evolving across metazoan lineages (Benoit et al., 2010; Kim et al., 2024; King & MacRae, 2015). Parallel expansions of SOD2 and SelR, both critical in mitigating damage from reactive oxygen species (ROS), indicate enhanced mitochondrial defenses in terrestrial lineages. SOD2, in particular, scavenges ROS generated during high metabolic activity (Bono, 2021; Kokoszka et al., 2001; Miao & St Clair, 2009; Y. Wang et al., 2018).

Parallelly evolving OGs were annotated to genes linked to key developmental signaling pathways, including the insulin/IGF signaling (IIS) pathway, a critical regulator of metabolism, growth, and reproduction. Duplications of insulin-like peptides (ILPs) and receptors (InRs) across arthropods suggest the evolutionary plasticity of this system (Nässel & Vanden Broeck, 2016). IIS has also been tied to reproductive strategies and environmental sensing, reinforcing its adaptive relevance during habitat transitions (Kremer et al., 2018; Wu & Brown, 2006). The expansion of IIS components in terrestrial arthropods may also have contributed to the evolution of more complex and plastic life histories, as this pathway integrates nutritional status, growth, reproduction, and environmental cues. Such functional versatility could have facilitated adaptive diversification during terrestrialization, allowing different lineages to fine-tune developmental timing, reproductive cycles, and metabolic trade-offs in response to variable terrestrial conditions.

Immune signaling pathways such as Toll, Imd, and JAK-STAT also showed either contraction or positive selection depending on the lineage (Amoyel & Bach, 2012; Zhou & Agaisse, 2012). The Toll and IMD pathways, central to bacterial and viral defense, show lineage-specific gene loss or redundancy compensation (O’Neal et al., 2023; Orús-Alcalde et al., 2021; Palmer & Jiggins, 2015). Additional pathways, including EGFR, Notch, Hedgehog, Wnt, and Hippo, are involved in morphogenesis, cell fate, and tissue organization and likely played key roles in enabling developmental flexibility and new body plans in terrestrial niches (Y. Chen et al., 2023; Li et al., 2022; Villarreal et al., 2015).

A prominent feature among expanded OGs is the regulatory complexity tied to gene expression modulation, including enriched GO terms for mRNA processing, alternative splicing, and transcription regulation. Regulatory adaptations such as alternative splicing have been documented as critical mechanisms in stress tolerance in extreme environments, as seen in *Belgica antarctica (Teets et al., 2012)* and *Birgus latro (Veldsman et al., 2021)*.

Moreover, the mixed patterns of expansion and contraction within gene families like CYPs, HSPs, and SLC transporters suggest a fine-scale evolutionary tuning, optimizing specific subfunctions while streamlining others. This balance reflects the complex interplay between innovation and efficiency, likely shaped by lineage-specific ecological constraints. Similar expansions of these gene families have also been reported across arthropods irrespective of habitat, as highlighted by Thomas et al. (2020), who documented widespread lineage-specific gains in categories such as HSPs, cytochrome P450s, and solute carrier (SLC) transporters. This indicates that terrestrialization likely built upon pre-existing genomic plasticity present within arthropods. Our contribution extends these observations by demonstrating that a subset of these families underwent parallel expansions specifically along terrestrial branches and under signatures of positive selection, distinguishing adaptive terrestrial shifts from general patterns of genomic diversification.

Together, these findings highlight that arthropod terrestrialization was not a product of isolated innovations but instead emerged through repeated, parallel genomic strategies that modified shared functional systems under convergent selective pressures. The interplay between gene family expansion and contraction, especially in pathways governing metabolism, stress tolerance, development, and immunity, reveals a deep evolutionary logic to how organisms repeatedly solve the challenges of colonizing land. The genomic parallelism observed across Arachnida, Myriapoda, and Hexapoda points to the emergence of a genomic toolkit for terrestrial adaptation—one that is flexible enough to accommodate lineage-specific ecological demands yet conserved in its core components. These results not only underscore the power of gene repertoire remodeling in enabling large-scale habitat transitions but also provide a valuable framework for identifying common molecular solutions to environmental pressures across Metazoa.

## Conclusions

Our study provides comprehensive genomic evidence that terrestrialization in arthropods was facilitated by parallel evolution of gene families under selection across independently evolved terrestrial lineages. By combining large-scale orthogroup reconstruction, gene family dynamics, and selection analysis, we identified a core set of gene families—related to cellular respiration, oxidative stress protection, transmembrane transport, moulting, and exoskeleton development—that likely form a shared genetic toolkit for terrestrial adaptation. The involvement of aquaporins, SODs, HSPs, CYPs, and SLC transporters underscores the importance of stress resilience and homeostasis in terrestrial environments. Moreover, the parallelism of changes in developmental and immune signaling pathways points to deep genomic remodeling in response to the ecological challenges of life on land.

Our findings contribute to the understanding of arthropod evolution and provide a genomic framework for exploring the genetic basis of terrestrial adaptation. The identification of specific gene families underpinning adaptation to new habitats highlights the intricate interplay between genomic changes and environmental challenges during the conquest of land by arthropods. These findings not only illuminate the molecular basis of arthropod terrestrialization but also offer a broader perspective on how parallelgenomic strategies enable complex habitat transitions across deep evolutionary time. While our binary classification of species as terrestrial or aquatic is appropriate for capturing the deep evolutionary transitions analyzed here, we acknowledge that this coarse ecological coding does not capture more recent or fine-scale habitat shifts (e.g., semi-terrestrial or amphibious lifestyles). As with any large-scale comparative genomic study, our conclusions are subject to several limitations. First, taxon sampling across arthropods remains uneven, reflecting biases in available genomic resources. Second, the functional annotations used for enrichment analyses are largely derived from model organisms and may overrepresent certain lineages, which can introduce interpretive uncertainty. Third, our analyses cannot fully disentangle terrestrialization from other correlated macroevolutionary processes such as diversification or body plan innovation. Addressing these challenges will require expanding genomic representation across transitional and dual-lifestyle taxa, incorporating transcriptomic and functional data, and applying more refined ecological classifications. Despite these caveats, the consistency of the observed patterns across independent lineages supports the robustness of our main conclusions and provides a strong foundation for future work on the genomic basis of ecological adaptation. Further studies leveraging these insights will deepen our understanding of evolutionary processes and biological innovations crucial for successful terrestrial colonization.

## Data Accessibility and Benefit-Sharing

Supplementary Material including support data and figures as well as scripts is available in GitHub (https://github.com/MetazoaPhylogenomicsLab/Benitez_Alvarez_2025_Arthropod_terrestrialization_MolEcol) and FigShare (https://figshare.com/articles/dataset/Parallel_genomic_remodeling_associated_with_independent_terrestrialization_events_in_arthropods/30428059).

## Acknowledgements

LA acknowledges funding from a Juan de la Cierva-Formación (grant agreement no. FJC2019-042184-I funded by MCIN/AEI/10.13039/501100011033). RF acknowledges support from the following sources of funding: Ramón y Cajal fellowship (grant agreement no. RYC2017-22492 funded by MCIN/AEI /10.13039/501100011033 and ESF ‘Investing in your future’), the Agencia Estatal de Investigación (project PID2019-108824GA-I00 funded by MCIN/AEI/10.13039/501100011033), the European Research Council (this project has received funding from the European Research Council (ERC) under the European’s Union’s Horizon 2020 research and innovation programme (grant agreement no. 948281)), the Human Frontier Science Program (grant no. RGY0056/2022) and the Secretaria d’Universitats i Recerca del Departament d’Economia i Coneixement de la Generalitat de Catalunya (AGAUR 2021-SGR00420). We also thank Centro de Supercomputación de Galicia and CSIC for access to computer resources (CESGA and DRAGO respectively).

## Author Contributions

VT, LA and RF designed the study. LB-A and VT performed the main analyses. VT and LA wrote the code and the pipeline to automate the methods. LA guided the analyses. All authors interpreted the results. LB-A, VT and RF wrote the first version of the manuscript. LA and RF supervised the study. RF provided resources. All authors revised and approved the final version of the manuscript.

